# Perfusion bioreactor culture of adipose-derived stromal cells on decellularized adipose tissue scaffolds enhances *in vivo* regeneration

**DOI:** 10.1101/2020.04.10.036194

**Authors:** Tim Tian Y. Han, Lauren E. Flynn

**Affiliations:** Department of Anatomy & Cell Biology, Schulich School of Medicine & Dentistry, The University of Western Ontario, London, Ontario, Canada, N6A 5C1; Department of Chemical and Biochemical Engineering, Thompson Engineering Building, The University of Western Ontario, London, Ontario, Canada, N6A 5B9; Bone and Joint Institute, The University of Western Ontario, London, Ontario, Canada, N6G 2V4

**Keywords:** adipose tissue engineering, adipose-derived stromal cells, bioreactors, bioscaffolds, decellularized adipose tissue, angiogenesis, hypoxic preconditioning

## Abstract

Adipose tissue engineering holds promise to address the unmet need in plastic and reconstructive surgery for strategies that promote the stable and predictable regeneration of adipose tissue for volume augmentation applications. Previous studies have demonstrated that decellularized adipose tissue (DAT) scaffolds can provide a pro-adipogenic microenvironment, and that seeding with adipose-derived stromal cells (ASCs) can enhance *in vivo* angiogenesis and adipogenesis within DAT implants. Recognizing that bioreactor systems can promote cell expansion and infiltration on tissue-engineered scaffolds, this study evaluated the effects of culturing human ASCs on DAT scaffolds within a perfusion bioreactor. Using this system, the impact of both shear stress stimulation and hypoxic preconditioning were explored *in vitro* and *in vivo*. Initial studies compared the effects of 14 days of culture within the perfusion bioreactor under 2% O_2_ or ~20% O_2_ on human ASC expansion and hypoxia inducible factor 1 alpha (HIF-1α) expression *in vitro* relative to static cultured controls. The findings indicated that culturing within the bioreactor under 2% O_2_ significantly increased ASC proliferation on the DAT, with a higher cell density observed in the scaffold periphery. HIF-1α expression was significantly higher when the scaffolds were cultured under 2% O_2_. Subsequent characterization in a subcutaneous implant model in athymic nude mice revealed that *in vivo* angiogenesis and adipogenesis were markedly enhanced when the ASCs were cultured on the DAT within the perfusion bioreactor under 2% O_2_ for 14 days prior to implantation relative to the other culture conditions, as well as additional freshly-seeded and unseeded DAT control groups. Overall, dynamic culture within the perfusion bioreactor system under hypoxia represents a promising approach for preconditioning ASCs on DAT scaffolds to enhance their capacity to stimulate blood vessel formation and infiltration, as well as host-derived adipose tissue regeneration.

## Introduction

The subcutaneous adipose tissue layer plays important structural and functional roles within the integumentary system^1^. Due to the limited capacity of adipose tissue for self-repair, damage or loss resulting from trauma, burns, tumour resections, disease or aging can alter the normal body contours and result in scarring, deformities and loss of function^2^. While reconstruction can have a major positive impact on self-image and quality of life, there are numerous limitations to the existing clinical interventions that rely on tissue transfer from other sites on the body or on current implants used for volume augmentation^3,4^.

Adipose tissue engineering represents a promising approach to address the unmet clinical need in plastic and reconstructive surgery for strategies that can promote the stable regeneration of healthy host-derived soft tissues^5,6^. Towards this goal, our lab has pioneered the development of off-the-shelf bioscaffolds derived from human decellularized adipose tissue (DAT), demonstrating that they provide a pro-adipogenic microenvironment both *in vitro* and *in vivo*^7,8^. DAT contains a diverse array of extracellular matrix (ECM) components, including a variety of matricellular proteins, small leucine-rich proteoglycans (SLRPs), growth factors and chemokines^9^ that may have bioactive effects on seeded and infiltrating host cell populations. Moreover, the DAT has similar biomechanical properties to native human fat^10,11^, which may be favorable for both implant integration and adipogenic differentiation^7,12^.

Previously, we demonstrated that seeding the DAT with allogeneic adipose-derived stromal cells (ASCs) enhanced angiogenesis and adipogenesis within the implants in an immunocompetent Wistar rat model^13^. More specifically, our study suggested that the ASCs indirectly contributed to host-derived adipose tissue regeneration through paracrine mechanisms, rather than through long-term engraftment and differentiation. It is well-established that ASCs secrete a variety of pro-regenerative growth factors and cytokines that can promote the recruitment of host progenitor cells, stimulate angiogenesis, and modulate the response of infiltrating immune cell populations, to help coordinate tissue regeneration^14–16^.

A key limitation to the previous approach was that the static seeding and culture methods employed resulted in low cellular infiltration and a heterogeneous distribution of the ASCs within the dense ECM of the DAT, which may have limited their long-term survival and functionality following *in vivo* implantation. As such, the primary goal of this study was to explore the potential of dynamic culture as a strategy to enhance human ASC expansion, infiltration and pro-regenerative function on the DAT bioscaffolds. In particular, perfusion bioreactors represent a promising approach for promoting cell growth on 3-D bioscaffolds by enhancing nutrient delivery and waste removal^17,18^. Further, the shear forces applied by these systems can modulate cell phenotype and function^19–22^, which may alter their capacity to stimulate regeneration.

In the current experimental design, human ASCs were pre-seeded on the DAT under dynamic conditions on an orbital shaker and then transferred into a custom-designed perfusion bioreactor system and cultured for 14 days. As oxygen tension has been shown to modulate ASC survival^23^, proliferation^24^, differentiation^25,26^, and pro-angiogenic factor production^14,27,28^ through the regulation of hypoxia inducible factor-1 alpha (HIF-1α)^29–31^, we explored the effects of culturing the cells under both normoxic (~20% O_2_) and hypoxic (2% O_2_) conditions. The selection of 2% O_2_ was based on studies showing enhanced ASC expansion *in vitro* at this level^25,32^. Initial studies focused on characterizing human ASC distribution, proliferation, and HIF-1α expression *in vitro* relative to control scaffold groups cultured under static conditions. Subsequently, *in vivo* testing was performed to probe the effects of dynamic culture within the perfusion bioreactor on human ASC retention, angiogenesis and adipogenesis within a subcutaneous implant model in athymic nude (*nu/nu*) mice relative to static cultured, freshly-seeded and unseeded DAT scaffold controls.

## Methods

### Materials

Unless otherwise stated, all chemicals and reagents were purchased from Sigma Aldrich Canada Ltd. (Oakville, Canada), and all antibodies were purchased from Abcam (Cambridge, USA).

### Adipose tissue processing

Subcutaneous adipose tissue was collected with informed consent from elective breast reduction or abdominoplasty surgeries at the University Hospital and St. Joseph’s Hospital in London, Canada. All studies were approved by the Human Research Ethics Board at Western University (HREB# 105426). The tissues were transported on ice in sterile phosphate buffered saline (PBS) with 2% bovine serum albumin (BSA) and processed within 2 h of collection for ASC isolation^33^ or DAT scaffold fabrication^7^ using published protocols.

ASCs were cultured in proliferation medium comprised of DMEM:Ham’s F12 supplemented with 10% fetal bovine serum (FBS) (Wisent, St. Bruno, Canada) and 100 U/mL penicillin/0.1 mg/mL streptomycin (1% pen-strep) (ThermoFisher, Waltham, USA) at 37°C with 5% CO_2_ and ~20% O_2_. Freshly isolated cells were passaged at 80% confluence, and passage 3 (P3) cells were used in all studies.

Following decellularization, the DAT was lyophilized and cut into scaffolds with a mass of 8 ± 1 mg. In preparation for seeding, the DAT scaffolds were rehydrated in deionized water for 24 h and then decontaminated by repeated rinsing in 70% ethanol. Next, the scaffolds were rinsed in sterile PBS and incubated in proliferation medium for 24 h.

### Scaffold seeding and bioreactor setup

To seed the scaffolds, 1 × 10^6^ P3 human ASCs were combined with individual DAT scaffolds in 3 mL of proliferation medium within 15 mL vented cap conical tubes (CellTreat Scientific Products, Pepperell, USA). The tubes were transferred into an incubator and agitated on an orbital shaker at a 15° incline and 100 RPM for 24 h (37°C, 5% CO_2_). Following seeding, the scaffolds were transferred into culture inserts that could be positioned within the individual sample chambers of the customized perfusion bioreactor system (Tissue Growth Technologies, Instron) (Supplemental Figure 1). The cell-seeded scaffolds were cultured under a flow rate of 0.5 mL/min for 14 days in proliferation medium under either normoxic (~20% O_2_: 5% CO_2_/95% air) or hypoxic (5% CO_2_/93% N_2_/2% O_2_) conditions, controlled through the use of a tri-gas incubator (ThermoFisher Forma Series II 3130), at 37°C. Static control scaffolds were included by culturing within the inserts under static conditions within vented cap conical tubes.

**Figure 1.**
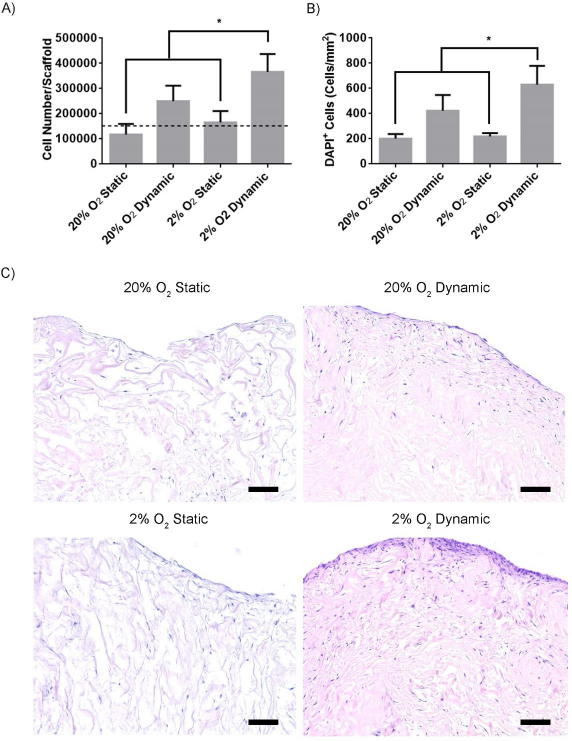
ASC density was enhanced on the DAT scaffolds following 14 days of culture within the perfusion bioreactor under 2% O_2_. A) Average number of ASCs/scaffold based on total dsDNA content following 14 days of culture. Dashed line represents the average cell density in the freshly-seeded group. Error bars represent standard deviation (n=4, N=3). B) The average number of ASCs per mm^2^ of the DAT bioscaffold as measured by DAPI quantification following 14 days of culture. Error bars represent standard deviation (n=3, N=3). * = 2% O_2_ dynamic is significantly different than 20% O_2_ static and 2% O_2_ static group (P<0.05). C) Representative H&E staining showing the distribution of ASCs on the DAT scaffolds following 14 days of culture. Scale bars represent 100 µm.

### Assessment of ASC density and distribution

After 14 days of culture, scaffold samples were collected (n=4 replicate scaffolds/trial, N=3 trials with different ASC donors) for analysis of total cell density using the Quant-iT^®^ PicoGreen^®^ dsDNA Assay Kit (ThermoFisher, Waltham, USA). For all trials, control groups were included of freshly-seeded (collected after the 24 h dynamic seeding phase) and unseeded DAT scaffolds. The scaffolds were finely minced with surgical scissors and digested in 1 mL of 4 mg/mL collagenase (Worthington Type I) in Kreb’s Ringer Buffer supplemented with 1% BSA, 2 mM glucose, and 25 mM HEPES under agitation at 100 RPM on an orbital shaker for 1 h at 37°C. The digested scaffolds were then centrifuged for 10 min at 1200 xg at room temperature, resuspended in Tris-EDTA buffer (10 mM Tris base, 1 mM EDTA, pH 8), and briefly sonicated using an ultrasonic dismembrator (ThermoFisher Model 100). The samples were then centrifuged at 5000 xg for 10 min at room temperature to remove debris, and the supernatant was analyzed using the PicoGreen^®^ Assay kit according to the manufacturer’s instructions using a CLARIOstar^®^ microplate reader (BMG Labtech, Guelph, Canada). All plates included a standard curve generated with known numbers of ASCs to estimate the total number of ASCs per scaffold.

To corroborate the PicoGreen^®^ data, the density of ASCs on the scaffolds was also assessed after 14 days of culture through quantification of DAPI staining (n=3, N=3). Scaffold samples were fixed in 4% paraformaldehyde for 24 h at 4°C before being embedded in paraffin and cut into 7 µm sections. Three non-adjacent cross-sections 100 µm apart were stained with DAPI, mounted and imaged with an EVOS^®^ FL Cell Imaging System (ThermoFisher) under the 20X objective. DAPI^+^ cells from 10 randomly selected non-adjacent fields from each scaffold cross-section were quantified using ImageJ software.

Hematoxylin & eosin (H&E) staining was also performed on scaffold cross-sections following standard procedures to assess the cell distribution within the DAT scaffolds after 14 days of culture (n=4, N=3). The samples were visualized using an EVOS^®^ XL Cell Imaging System (ThermoFisher).

### Immunohistochemical analysis of Ki-67 and HIF-1α expression

Immunohistochemical staining was performed on additional scaffold sections (3 non-adjacent sections at least 100 µm apart/scaffold; n=3, N=4) to examine the expression of Ki-67 as a proliferation marker and HIF-1α to probe potential changes under hypoxia. Heat-mediated antigen retrieval was performed in target antigen retrieval solution (Agilent, Santa Clara, USA) for 25 min. The slides were allowed to cool for 25 min and then washed in PBS and blocked in PBS-T (0.1% Tween-20) supplemented with 10% goat serum for 1 h at room temperature. Sections were then incubated with rabbit anti-Ki-67 antibody (1:100 dilution) or mouse anti-HIF-1α antibody (Novus Biologicals, Oakville, Canada; 1:100 dilution) overnight at 4°C. Alexa Fluor^®^ 594 conjugated goat anti-rabbit (1:100 dilution) or Alexa Fluor^®^ 680 conjugated goat anti-mouse (1:100 dilution) secondary antibody was subsequently applied for 1 h at room temperature and the samples were mounted in Fluoroshield Mounting Medium with DAPI (Abcam, Cambridge, USA). Stained sections were visualized with the EVOS^®^ FL Cell Imaging System, and 10 randomly selected non-adjacent fields of view from both the peripheral (< 200 µm from the scaffold edge) and central (>200 µm from the scaffold edge) regions of the scaffolds were imaged under the 20X objective. The number of Ki-67^+^DAPI^+^ cells and HIF-1α^+^DAPI^+^ within each region were quantified using ImageJ Software.

### Subcutaneous implantation surgeries

Following the *in vitro* testing, studies were performed to assess the effects of bioreactor culture on the capacity of the human ASCs to stimulate *in vivo* angiogenesis and adipogenesis within the DAT scaffolds. More specifically, these studies compared 6 scaffold groups including the ASC-seeded scaffolds that had been cultured for 14 days within the perfusion bioreactor or under static conditions at both 20% and 2% O_2_ (i.e. (i) 20% O_2_ static, (ii) 20% O_2_ dynamic, (iii) 2% O_2_ static, (iv) 2% O_2_ dynamic) to (v) freshly-seeded (after the 24 h seeding phase) and (vi) unseeded DAT scaffold controls. For each scaffold group, duplicate samples were implanted for each trial and a total of 3 trials were performed using ASCs isolated from 3 different donors. All studies followed the Canadian Council on Animal Care (CCAC) guidelines and the protocols were approved by the Animal Care Committee at Western University (Protocol # 2015-049).

The scaffolds were implanted subcutaneously in female Nu-*Foxn1^nu^* mice (Charles River Laboratories, Sherbrooke, Canada) that were 10-13 weeks of age for 4 and 8 weeks. A total of 36 mice were used for this study (N=6 mice per scaffold group at each timepoint). The mice were anaesthetized with isoflurane and given pre-operative analgesics via subcutaneous injections of meloxicam (2 mg/kg loading dose; 1 mg/kg follow up dose at 24 h) and bupivacaine (2 mg/kg). Two small incisions (~7 mm) were made on the dorsa of each mouse below each scapula, and pockets were created using blunt-ended forceps. One scaffold was then carefully positioned within each pocket and the incisions were closed with surgical staples. At each timepoint, the mice were euthanized by CO_2_ overdose and the scaffolds were excised within their surrounding tissues. The samples were fixed in 4% paraformaldehyde overnight at 4°C, prior to paraffin embedding and sectioning (7 µm thick) in preparation for histological and immunohistochemical analyses. For all staining analyses, 3 cross-sections per implant (N=6 per time point) were analyzed that were a minimum of 100 µm apart.

### Masson’s trichrome staining

Masson’s trichrome staining was performed using standard procedures to assess implant integration and remodeling, as well as the density of erythrocyte-containing blood vessels within the scaffolds. The entire scaffold cross-section in each sample was imaged using the EVOS^®^ XL Cell Imaging System at 20X magnification and combined into a single cross-sectional image using Adobe^®^ Photoshop^®^ CS6 Software. Within each complete cross-section, the number of erythrocyte-containing blood vessels was quantified within the DAT implant to determine the overall blood vessel density. Further, the distance to the closest scaffold edge and diameter were measured for each vessel using ImageJ Software. Vessel infiltration within 500 µm of the scaffold surface was analyzed using frequency distribution graphs with a bin size of 25 µm.

### Immunohistochemical analysis of perilipin expression

At both timepoints, immunohistochemical staining was performed to assess perilipin expression as a marker of adipocyte formation within the DAT implants. The sections were prepared as described above and incubated overnight in rabbit anti-perilipin A antibody (1:200 dilution) at 4°C, followed by a goat anti-rabbit HRP-conjugated secondary antibody (1:500 dilution) for 1 h at room temperature. The samples were then processed with a DAB peroxidase (HRP) substrate kit (Vector Labs, Burlington, Canada), followed by counterstaining with hematoxylin and mounting with Permount^®^ Mounting Medium (ThermoFisher, Waltham, USA). The entire scaffold region was imaged at 10X magnification and stitched into a single cross-sectional image as previously described. Positive pixel counting was performed to analyze the relative perilipin expression levels between the implant groups, with the values normalized to the total scaffold area within each cross-section measured using ImageJ Software.

### Immunohistochemical analysis of human ASC engraftment and differentiation

To detect the human ASCs within the DAT implants, cross-sections were stained for human-specific Ku80^34^ using a rabbit anti-Ku80 antibody (Cedarlane, Burlington, Canada; 1:100 dilution) followed by Alexa Fluor^®^ 594 conjugated goat anti-rabbit (1:200 dilution) secondary antibody as described above. The stained cross-sections were imaged using the EVOS^®^ FL Cell Imaging System. Quantification of Ku80^+^DAPI^+^ cells was performed on 10 randomly selected non-overlapping fields of view taken at 20X magnification using ImageJ analysis software.

To probe whether the human ASCs were differentiating into adipocytes within the DAT implant region, co-staining was performed using rabbit anti-perilipin A antibody as an adipocyte marker in combination with a mouse anti-human mitochondria (hMito) antibody conjugated with Cy3 (EMD Millipore, Burlington, USA; 1:100 dilution). hMito was used as an alternative to Ku80 based on the availability of antibodies compatible for co-staining. The samples were incubated overnight at 4°C and then incubated for 1 h at room temperature with a goat anti-rabbit secondary antibody conjugated with Alexa Fluor^®^ 680 (1:100 dilution). The stained sections were visualized with the EVOS^®^ FL Cell Imaging System and imaged under 20X magnification.

### Statistical analysis

All numerical values are represented as the mean ± standard deviation. Statistical analyses were performed using GraphPad Prism^®^ version 6 by one way or two-way ANOVA and followed by a Tukey’s post-hoc test. Differences were considered statistically significant at p<0.05.

## Results

### Dynamic culturing in the bioreactor under hypoxic conditions for 14 days enhanced ASC density on the DAT bioscaffolds

To quantitatively assess cell density on the DAT, the scaffolds were digested and analyzed with the PicoGreen^®^ dsDNA Assay. Freshly-seeded scaffolds had an average of 1.49 ± 0.19 × 10^5^ cells per scaffold following the 24 h seeding period. After 14 days of culture, the total cell density was significantly higher in the 2% O_2_ dynamic group as compared to the freshly-seeded scaffolds and the 20% O_2_ static and 2% O_2_ static groups (Figure 1A). DAPI staining showed similar patterns in terms of the ASC density on the DAT scaffolds, corroborating the PicoGreen^®^ results (Figure 1B).

To complement the quantitative findings, H&E staining was performed to assess the ASC distribution within the DAT scaffolds following the 14-day culture period. The ASCs were predominantly localized to the surface of the DAT scaffolds in the samples that were cultured under static conditions at both 20% and 2% O_2_ (Figure 1C). A higher density of ASCs was observed in the scaffolds cultured for 14 days within the bioreactor, with a dense cell layer observed along the surface of the scaffolds. Consistent with the PicoGreen^®^ results, the 2% O_2_ dynamic group appeared to have a qualitatively higher density of cells, with greater infiltration into the central scaffold regions.

### Dynamic culturing in the bioreactor under hypoxic conditions for 14 days enhanced Ki-67 expression in the scaffold periphery

Immunostaining identified a low density of Ki-67^+^DAPI^+^ cells distributed throughout the scaffolds at 14 days. Qualitatively, there appeared to be more Ki-67^+^DAPI^+^ cells in the 2% O_2_ dynamic group, with a higher density in the scaffold periphery (Figure 2A). Representative images of the central scaffold regions are shown in Supplemental Figure 2. These observations were confirmed by quantification of the number of Ki-67^+^DAPI^+^ cells within the peripheral (< 200 µm from edge) and central (> 200 µm from edge) regions of the scaffolds, which revealed that there was a significantly higher density of Ki-67^+^DAPI^+^ cells in the periphery in the 2% O_2_ dynamic group as compared to both the peripheral and central regions of all other culture conditions. Additionally, a greater density of Ki-67^+^DAPI^+^ cells was measured in the central region of 2% O_2_ dynamic group as compared to the central region of 2% O_2_ static group (Figure 2B).

**Figure 2.**
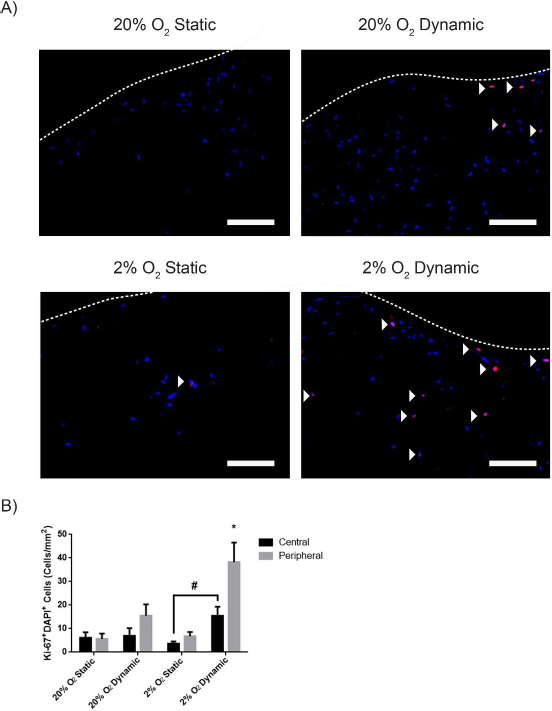
Ki-67 expression at 14 days was enhanced in the ASCs dynamically cultured on the DAT scaffolds within the perfusion bioreactor under 2% O_2_. A) Representative images showing Ki-67^+^ cells (red) counterstained with DAPI (blue) in the peripheral regions of the scaffolds after 14 days of culture. White arrowheads denote positively labelled cells and dashed lines represents the edge of the bioscaffolds. Scale bars represent 100 µm. B) The average number of Ki-67^+^DAPI^+^ cells in the peripheral and central regions of the scaffolds at 14 days. Error bars represent standard deviation (n=3, N=4). * = significant difference between 2% O_2_ dynamic group and all other groups, # = significant difference between the central region of 2% O_2_ dynamic and 2% O_2_ static groups (p<0.05).

### Culturing under hypoxia increased the number of HIF-1α^+^ cells within the scaffolds at 14 days

Immunohistochemical staining revealed that there were very few HIF-1α^+^DAPI^+^ cells in both the peripheral (< 200 µm from edge) and central (> 200 µm from edge) regions of the scaffolds cultured either dynamically or statically for 14 days under 20% O_2_ (Figure 3A). In contrast, HIF-1α expression was observed in the majority of the cells in the 2% O_2_ static and dynamic groups. Representative images of the central scaffold regions are shown in Supplemental Figure 3. Quantification of the staining indicated that the density of HIF-α^+^DAPI^+^ cells was significantly higher in the 2% O_2_ dynamic group as compared to all other groups within both the peripheral and central scaffold regions (Figure 3B). In addition, the density of HIF-1α^+^ DAPI^+^ cells was also significantly greater in the 2% O_2_ static group as compared to the 20% O_2_ static and dynamic groups within both the peripheral and central scaffold regions (Figure 3B).

**Figure 3.**
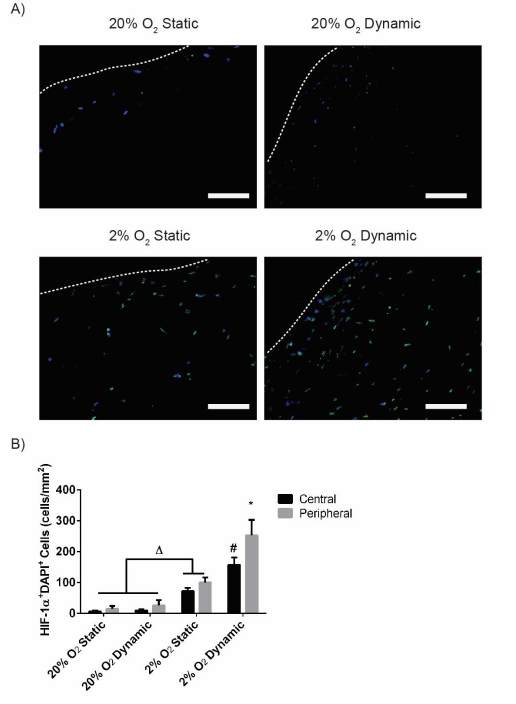
HIF-1α expression was enhanced at 14 days in the ASCs cultured on the DAT scaffolds under 2% O_2_. A) Representative immunostaining showing HIF-1α^+^ ASCs (green) counterstained with DAPI (blue) in the peripheral (< 200 µm from the edge) regions of the scaffolds following 14 days of culture. Scale bars represent 100 µm. Dashed lines represent the border of the bioscaffolds. B) The average number of HIF-1α^+^DAPI^+^ cells in the peripheral and central regions of the scaffolds at 14 days. Error bars represent standard deviation (n=3, N=4). * = significant difference between the peripheral region of the 2% O_2_ dynamic group and both the central and peripheral regions of all other groups, # = significant difference between the central region of the 2% O_2_ dynamic group and the central region of all other conditions, Δ = significant difference between the 2% O_2_ static group as compared to the 20% O_2_ static and 20% O_2_ dynamic groups in both regions (P<0.05).

### DAT scaffolds cultured within the bioreactor for 14 days under hypoxic conditions showed enhanced blood vessel formation at 4 and 8 weeks following subcutaneous implantation

*In vivo* studies were subsequently performed using a subcutaneous implantation model in *nu/nu* mice to compare angiogenesis and adipogenesis in the scaffolds that had been cultured for 14 days within the bioreactor or statically (20% O_2_ static, 20% O_2_ dynamic, 2% O_2_ static, 2% O_2_ dynamic) to freshly-seeded and unseeded controls. Masson’s trichrome staining at 4 and 8 weeks revealed that the scaffolds were well integrated into the host tissues and showed evidence of implant remodeling (Figure 4A). Small blood vessels and adipocytes were observed in the peripheral regions of the DAT implants starting at 4 weeks. At 8 weeks, more vessels were observed in the 2% O_2_ dynamic group as compared to all other conditions, along with a greater number of adipocytes extending into the more central scaffold regions (Figure 4A).

**Figure 4.**
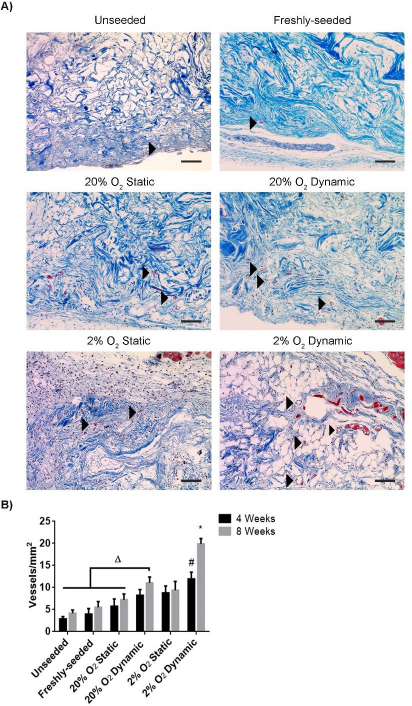
The density of erythrocyte-containing blood vessels was enhanced at 4 and 8 weeks in the DAT scaffolds that were cultured in the bioreactor under 2% O_2_ for 14 days prior to subcutaneous implantation in *nu/nu* mice. A) Representative Masson’s trichrome staining of the DAT implants at 8 weeks, with black arrowheads highlighting erythrocyte-containing blood vessels within the scaffolds. Scale bars represent 100 µm. B) Density of erythrocyte-containing blood vessels within the scaffolds at 4 and 8 weeks. Error bars represent standard deviation (n=3 cross-sections/implant, N=6 implants per group).* = 2% O_2_ dynamic group at 8 weeks is significantly different compared to all groups at both 4 and 8 weeks, # = significant difference between 2% O_2_ dynamic group and all other groups at 4 weeks, Δ = 20% O_2_ dynamic group is significantly different than the indicated groups at both 4 and 8 weeks (p<0.05).

Quantification of the number of erythrocyte-containing blood vessels in the scaffolds revealed that there was a significantly higher vessel density in the 2% O_2_ dynamic group as compared to all other conditions at both 4 and 8 weeks (Figure 4B), with a significant difference between the two timepoints for this group. Further, the vessel density in the 20% O_2_ dynamic group at 8 weeks was significantly greater than the unseeded, freshly-seeded and 20% O_2_ static groups at both 4 and 8 weeks. The diameter of the erythrocyte-containing blood vessels was also measured, with results indicating that there were no significant differences in the average diameter between the groups at both 4 and 8 weeks, with an average diameter in the range of ~25 - 35 µm (Supplemental Figure 4A).

Blood vessel infiltration was also probed by measuring the shortest distance from each vessel to the scaffold periphery. No significant difference in the average distance of blood vessel infiltration was measured between the groups at 4 weeks, with the average values ranging from ~70 - 125 µm and a high degree of variability observed (Supplemental Figure 4B). At 8 weeks, the average distance of vessel infiltration was significantly higher in the 2% O_2_ dynamic group (198.8 ± 32.1 µm) as compared to all other groups with the exception of the 20% O_2_ static (100.7 ± 38.3 µm) and 20% O_2_ dynamic (125.1 ± 46.6 µm) groups.

Vessel infiltration within 500 µm of the scaffold surface was further analyzed by frequency distribution plots with pooled counts of the total number of vessels within 25 µm intervals for each of the scaffold groups (Figure 5). Based on this analysis, an increase in the number of vessels within the first 100 µm of the scaffolds was observed in the 2% O_2_ dynamic and static groups at 4 weeks, suggesting that hypoxic culture promoted vessel formation in the scaffold periphery. At 8 weeks, a shift towards greater infiltration (i.e. rightwards) was observed for both the 2% O_2_ dynamic and 20% O_2_ dynamic groups, suggesting that dynamic culture enhanced the number of vessels infiltrating into the 100 – 200 µm depth range, with a higher number of vessels in the 2% O_2_ dynamic group in this range, as well as farther into the scaffolds. Interestingly, the distributions for the freshly-seeded and unseeded groups were similar at both timepoints, indicating that the ASCs had little impact on vessel formation within the DAT under the conditions in this study. While the distribution for the 20% O_2_ static group was similar to these groups at 4 weeks, an increase in the number of vessels within the first 50 µm was observed in this group at 8 weeks, suggesting that there was some benefit of culturing the ASCs on the scaffolds prior to implantation in terms of vessel formation within the most peripheral regions of the DAT scaffolds (Figure 5).

**Figure 5.**
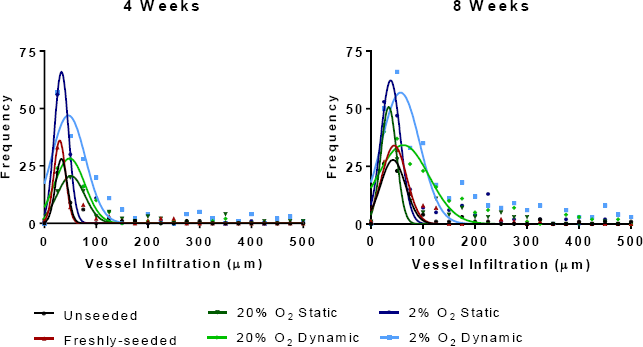
Culturing the ASCs on the DAT bioscaffolds modulated blood vessel infiltration into the DAT implants over time. Frequency distribution plots showing the total number of erythrocyte-containing blood vessels at increasing depths from the scaffold periphery within the DAT implants. Coloured lines represent non-linear Gaussian fits of the data sets. Pooled data is shown representing the combined counts from 3 cross-sections at varying depths from 6 implants per condition at both 4 and 8 weeks.

### DAT scaffolds cultured within the bioreactor for 14 days under hypoxic conditions showed enhanced adipose tissue remodeling at 8 weeks following subcutaneous implantation

Immunohistochemical staining for perilipin was performed to identify adipocytes containing intracellular lipid as a measure of adipose tissue remodeling within the implanted scaffolds at 4 (Supplemental Figure 5) and 8 weeks (Figure 6A). At 4 weeks, a small number of adipocytes were observed within the peripheral regions of the scaffolds in all of the groups. At 8 weeks, there were qualitatively more adipocytes in the scaffolds that incorporated ASCs as compared to the earlier timepoint, with more perilipin^+^ cells observed in the 2% O_2_ dynamic group as compared to all other conditions.

**Figure 6.**
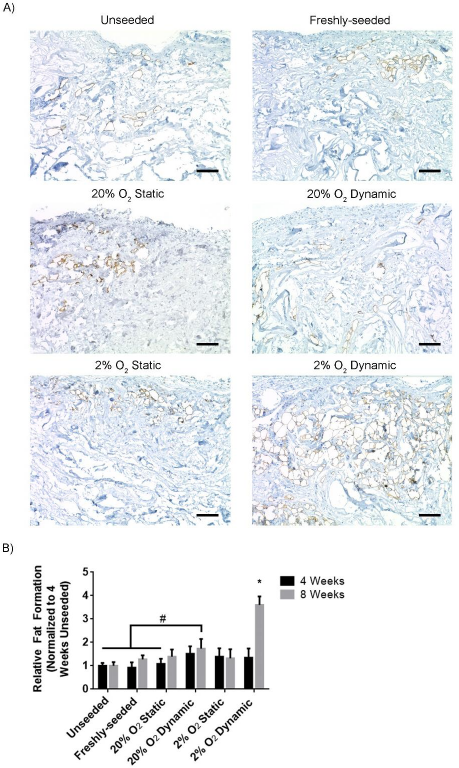
Remodeling of the DAT into adipose tissue was enhanced in the 2% O_2_ dynamic group at 8 weeks following subcutaneous implantation in the *nu/nu* mouse model. A) Representative immunostaining showing regions of perilipin^+^ adipocytes (brown) within the implanted scaffolds at 8 weeks. Scale bars represent 100 µm. B) The expression of perilipin relative to the unseeded group at 4 weeks. Error bars represent standard deviation (n=3 cross-sections/implant, N=6 implants per group). * = significant difference between the 2% O_2_ dynamic group at 8 weeks and all other groups at both 4 and 8 weeks, as well as the 2% O_2_ dynamic group at 4 weeks, # = significant difference between the 20% O_2_ dynamic group at 8 weeks and the indicated groups (P<0.05).

Semi-quantitative image analysis confirmed that the relative levels of perilipin expression within the implant region were similar between the groups at 4 weeks (Figure 6B). In contrast, at 8 weeks, the 2% O_2_ dynamic group showed significantly greater perilipin expression levels as compared to all other groups at both 4 and 8 weeks, with a significant increase observed within this group from 4 to 8 weeks. Collectively, these findings are indicative of enhanced remodeling of the DAT implant into adipose tissue in the 2% O_2_ dynamic condition. Further, perilipin expression was significantly higher in the 20% O_2_ dynamic group relative to the unseeded and freshly-seeded groups at both 4 and 8 weeks, as well as the 2% O_2_ static group at 4 weeks (Figure 6B).

### Human ASCs were detected in the DAT implants at similar levels at both 4 and 8 weeks

Immunohistochemical staining was performed for human-specific Ku80 to identify human ASCs within the DAT implants at both 4 (Supplemental Figure 6) and 8 weeks (Figure 7A). Ku80^+^ cells were visualized throughout the seeded scaffolds at both timepoints. Quantification of the number of Ku80^+^DAPI^+^ cells within the scaffold regions revealed that there were no significant differences in the density of human ASCs detected between the seeded implant groups at both 4 and 8 weeks (Figure 7B).

**Figure 7.**
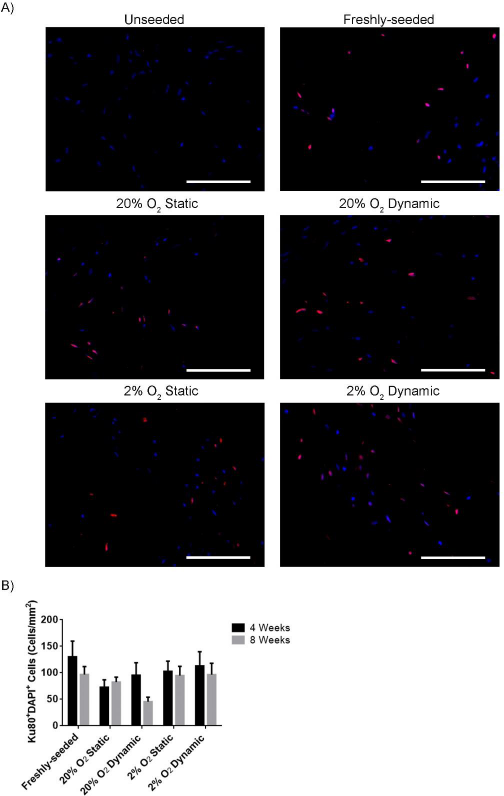
Ku80^+^ cell density was similar between all seeded groups at both 4 and 8 weeks post-implantation. A) Representative immunostaining showing Ku80 labelled cells (red) counterstained with DAPI (blue). Scale bars represent 100 µm. B) Bar graph showing the density of Ku80^+^DAPI^+^ ASCs in the implants at 4 and 8 weeks. Data was compiled from 6 implants per group (N=6), with 3 cross-sections per scaffold taken from varying depths and 10 random fields analyzed per cross-section. Error bars represent standard deviation.

### No evidence of human ASC differentiation into adipocytes in vivo

Co-staining with a human anti-mitochondrial (hMito) antibody in combination with perilipin was performed to probe human ASC differentiation into adipocytes within the DAT implants. No perilipin^+^hMito^+^ cells were identified, indicating that the newly-forming adipocytes within the DAT implants were predominantly host-derived (Figure 8).

**Figure 8.**
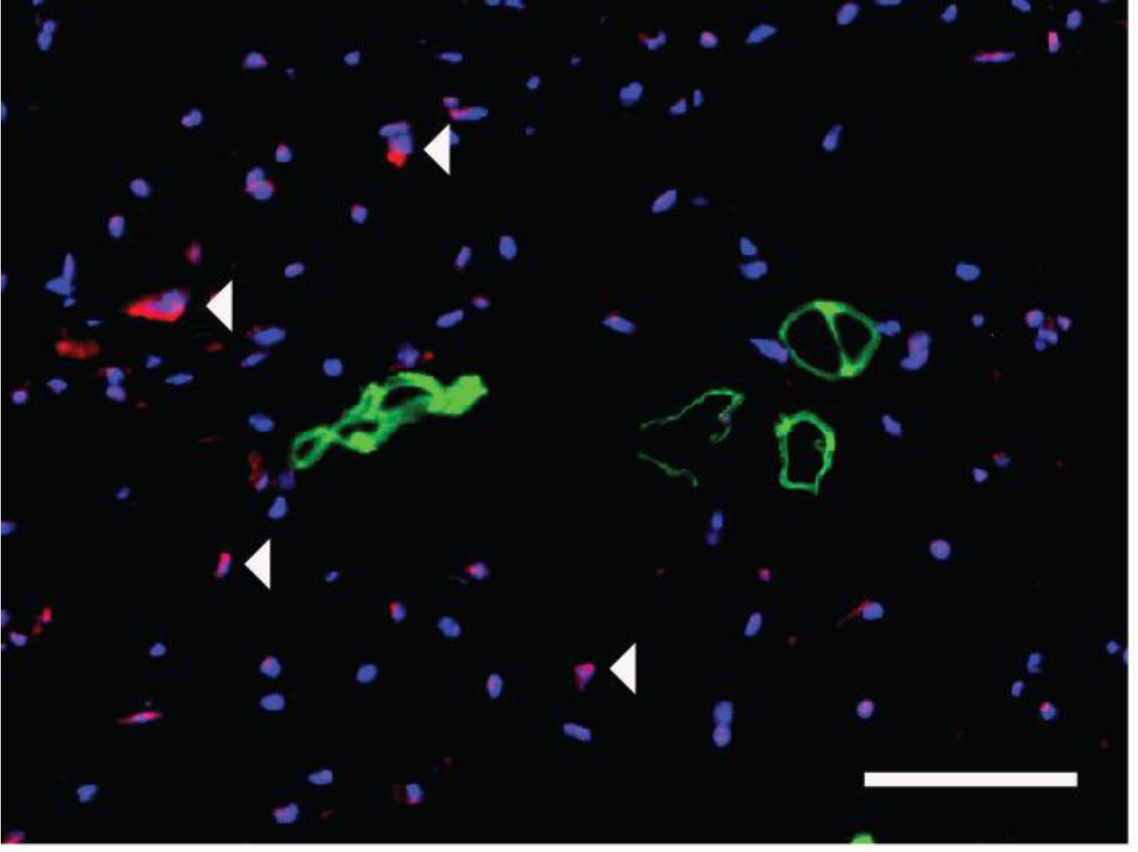
Staining patterns suggest that the human ASCs did not differentiate into adipocytes within the DAT implants over 8 weeks. Representative image showing distinct hMito^+^ (red) and perilipin^+^ (green) cell populations with DAPI (blue) counterstaining. White arrowheads indicate human ASCs in close proximity to perilipin^+^ cells. Scale bars represent 50 µm.

## Discussion

New strategies that can promote the regeneration of healthy host-derived adipose tissue and enable stable and predictable volume augmentation would be of great value for a broad range of applications in plastic and reconstructive surgery. With the long-term goal of advancing towards clinical translation, our lab has been developing a broad range of off-the-shelf bioscaffolds derived from the ECM of human adipose tissue^7,8,35–37^. Our previous *in vivo* study exploring DAT scaffolds in a subcutaneous implant model in immunocompetent rats indicated that seeding the DAT with allogeneic rat ASCs significantly enhanced blood vessel formation and remodeling of the DAT into host-derived adipose tissue^13^. In that study, the DAT scaffolds were seeded with the ASCs within tissue culture inserts and then cultured statically for 72 h to promote cell attachment. However, these static seeding and culture methods result in a heterogeneous and sparse distribution of ASCs on the relatively dense intact DAT scaffolds, with limited cell infiltration into the central scaffold regions. We postulated that having a higher density and more even distribution of cells throughout the scaffold may be favorable for enhancing the rate and extent of adipose tissue regeneration within the DAT. Further, in the current study, we also extended from our previous *in vivo* work by investigating human ASCs as a more clinically-relevant cell population.

Dynamic culture methods can enhance cell proliferation and infiltration into 3-D porous scaffolds by promoting the exchange of nutrients and waste, as well as by providing mechanical stimulation that can modulate cell function^38–40^. Stirred culture systems, such as spinner flasks, have been most extensively explored for the expansion of mesenchymal stromal cell (MSC) populations including ASCs, incorporating a range of microcarrier types to support cell attachment and growth^41–47^. Using an alternative design, Papadimitropoulos *et al.* investigated the use of a ceramic scaffold-based perfusion bioreactor for human bone marrow-derived MSC expansion^48^, and Kim and Ma applied a perfusion bioreactor system for the expansion of human MSCs from bone marrow on polyethylene terephthalate (PET) scaffolds^49^. In general, the overall goal in these strategies was to harness the benefits of dynamic culture within a 3-D system to better preserve MSC function in terms of their capacity to proliferate and differentiate towards the adipogenic, osteogenic and/or chondrogenic lineages, as compared to the progressive functional decline that is observed when the cells are expanded on 2-D tissue culture polystyrene under static conditions^44,50^. The cells expanded within these systems can typically be released, such as through enzymatic digestion, and subsequently incorporated within a range of delivery platforms depending on the application of interest^51,52^.

In the current study, an integrated approach was applied using a custom 3-D perfusion bioreactor to expand the human ASCs on the same DAT scaffold on which they would be delivered. While similar perfusion bioreactor strategies have been explored for bone tissue engineering applications using biomaterials with properties that mimic bone^51,52^, to the best of our knowledge this is the first study to investigate the use of a scaffold-based perfusion bioreactor for adipose tissue regeneration. Culturing within the perfusion bioreactor system was shown to enhance the cell density within the peripheral regions of the DAT scaffolds, with a more marked effect in the samples that were cultured under 2% O_2_. These findings supporting the benefits of perfusion culture are similar to a previous study using a double chamber perfusion bioreactor that demonstrated that media perfusion enhanced bone marrow-dervied MSC homogeneity within decelluarized porcine tracheal scaffolds^53^. Additionally, a study by Sailon *et al.* showed that that the density of MC3T3-E1 murine pre-osteoblastic cells was enhanced on porous polyurethane scaffolds when cultured within a perfusion bioreactor system as compared to static controls, with the cells typically localized in the more porous regions of the scaffolds^54^. The enhanced cell density within the peripheral regions of the DAT suggests that there may have been limited media perfusion into the central regions of the scaffolds over time, which is a potential limitation. Regardless, the *in vivo* findings clearly support that culturing the ASCs on the DAT within the perfusion bioreactor under 2% O_2_ significantly augmented angiogenesis and adipogenesis within the implants, supporting the benefits of the approach.

Based on the PicoGreen and Ki-67 staining results, culturing under dynamic conditions within the bioreactor stimulated ASC proliferation on the DAT. Interestingly, there was no benefit of culturing the scaffolds under static conditions for 14 days in terms of the total cell number per scaffold relative to the freshly-seeded group. In addition to promoting ASC proliferation, bioreactor culture may have enhanced long-term cell survival^55,56^, contributing to the higher cell density observed. Further, the media used in this study contains both mitotic factors as well as potent chemotactic factors^57,58^ that may have promoted cell migration from the scaffold interior to the more highly perfused periphery.

The ASCs cultured both statically and dynamically under 2% O_2_ were also shown to express HIF-1α, consistent with hypoxia in adipose tissue being < 5% O_2_^59,60^. HIF-1α activation is associated with MSC proliferation^61,62^, which may have contributed to the enhanced Ki-67 expression in the hypoxic dynamic group. A previous study by Choi *et al.* demonstrated that culturing human ASCs under 2% O_2_ increased proliferation relative to culture under ~21% O_2_ through HIF-1α-mediated mechanisms^26^. In a more recent study, human ASCs were found to be more proliferative when cultured under 1% O_2_ as compared to 2% O_2_ for 7 days^31^. Moreover, the increased proliferation was shown to be mediated through the HIF-1α-dependent upregulation of bFGF^31^. In addition to stimulating proliferation, HIF-1α is an important transcription factor that regulates the expression of a range of angiogenic growth factors^63^. As such, it is possible that culturing under 2% O_2_ may have modulated the paracrine factor secretion profile of the ASCs within the DAT, which would be interesting to explore in future work.

Following *in vitro* characterization, *in vivo* testing was performed to assess the effects of bioreactor culture on the capacity of the ASCs to promote angiogenesis and adipogenesis within the DAT. Athymic nude mice, which have been used in a number of adipose tissue engineering studies^64–71^, were applied to enable the investigation of human ASCs. Angiogenesis was probed by quantifying the number of erythrocyte-containing blood vessels within the implants, indicative of functional vasculature. Previously in the rat model, we found that seeding the DAT with allogeneic rat ASCs resulted in an increase in the number and diameter of erythrocyte-containing blood vessels within the implants starting at 8 weeks^13^. In the current study, the vessel density was significantly higher in the 2% O_2_ dynamic group at both 4 and 8 weeks as compared to all other conditions. Since the ASC density at 14 days was significantly higher in this group, it is possible that enhanced cell delivery may have contributed to increased angiogenesis. A study by Al-khaldi *et al.* demonstrated there was a dose-dependent increase in *in vivo* blood vessel formation at 28 days when varying densities of mouse MSCs (up to 2 × 10^6^ cells/mL) were delivered subcutaneously in Matrigel in C57BL/6 mice^72^. In addition to vessel density, the increased blood vessel infiltration observed in the hypoxic bioreactor group could be an indicator of an enhanced vessel sprouting effect. Previous studies have shown that VEGF and bFGF are key paracrine factors that are crucial in driving vessel sprouting^27^, which are secreted by ASCs at enhanced levels under hypoxia^29,30^. Moreover, there is also evidence to support that MSCs subjected to shear stress induced by dynamic culturing systems can show enhanced VEGF secretion^19,21^. As discussed, it would be worthwhile to explore the effects of dynamic culture on the DAT on ASC phenotype and paracrine function in future studies.

In addition to enhanced angiogenesis, culturing the ASCs on the DAT within the bioreactor under 2% O_2_ for 14 days prior to implantation resulted in a significant increase in the number of perilipin^+^ adipocytes within the implants at 8 weeks as compared to all other groups, consistent with enhanced remodeling of the DAT into adipose tissue. Adipose tissue regeneration is typically a gradual process that is coupled with angiogenesis, as a rich vascular supply is required to support mature adipocytes^73,74^. In our previous rat study, we found that seeding the DAT with allogeneic rat ASCs enhanced adipogenesis within the implants at both 8 and 12 weeks^13^. However, the response at 8 weeks was quite variable, with 20.1 ± 15.3% of the seeded DAT having remodeled into mature fat, as compared to 3.9 ± 5.7% in the unseeded group, based on area analysis of Masson’s trichrome stained cross-sections^13^. The lack of difference observed between the other ASC-seeded groups in the current study relative to the unseeded controls may be related to species variability in terms of both the ASC source and animal model employed, as well as to differences in the ASC dose delivered. In addition, it is possible that further differences would be observed between the groups at later timepoints. Regardless, our findings clearly demonstrate that dynamic culture of human ASCs on the DAT within the perfusion bioreactor under 2% O_2_ significantly augmented both angiogenesis and adipogenesis within the implants at 8 weeks, supporting the use of this new strategy for adipose tissue regeneration.

While the density of ASCs was higher on the 2% O_2_ dynamic scaffolds at the time of implantation, there were no significant differences between the groups in terms of the number of human ASCs detected within the implants at both 4 and 8 weeks. It is possible that the enhanced vascularization observed in the bioreactor-cultured samples may have contributed to this decline by promoting the recruitment of immune cell populations such as monocytes/macrophages that may recognize the human ASCs as being foreign. In our previous study, using fluorescence *in situ* hybridization (FISH) to track the allogeneic ASCs sourced from male donors via the y-chromosome, a progressive reduction in the number of donor cells within the DAT implants was observed over time, with only a small number of donor ASCs visualized at 8 weeks and no positive cells detected at 12 weeks, indicating that the newly-formed adipose tissues were predominantly host-derived^13^. Similarly, Parisi-Amon *et al.* showed that mouse ASCs delivered subcutaneously in nude mice within a peptide-based hydrogel showed an 80% reduction in exogenous cells after 7 days *in vivo* based on bioluminescence imaging^75^. Additionally, a more recent study using a poly(N-isopropylacrylamide)-polyethylene glycol co-polymer to encapsulate human ASCs demonstrated ~13% retention at 14 days following subcutaneous delivery in athymic nude mice^76^. The persistence of a relatively high density of human ASCs in the current study at 8 weeks in all of the seeded groups suggests that the DAT may provide a supportive platform for their delivery. The functional role of these cells remains unclear, but the staining patterns suggest that there was limited differentiation into adipocytes. It would be interesting to explore how long these cells persist in future studies, as well as probing differences in possible paracrine-mediated effects at later timepoints including host adipogenic progenitor cell recruitment.

## Conclusions

The perfusion bioreactor system represents a novel and promising platform for the dynamic expansion of pro-regenerative cell populations on 3-D bioscaffolds for adipose tissue engineering, and also an advancement towards a clinically viable strategy to support adipose tissue regeneration. The findings in this study demonstrated that the perfusion bioreactor system is effective at increasing the density of ASCs on the DAT, which suggests that it could address the previous limitations of the ASC-seeded DAT bioscaffolds. Increasing the ASC density on the DAT bioscaffolds via culturing within the perfusion bioreactor system under hypoxia subsequently translated into an enhanced capacity to support blood vessel formation and adipose tissue remodelling *in vivo*. Collectively, the body of work in the present study provided support for the use of a perfusion bioreactor system to increase the pro-regenerative potential of the ASC-seeded bioscaffolds. Moving forward, future studies should explore the mechanisms through which the ASC-seeded DAT supports angiogenesis and adipogenesis *in vivo* as the next step towards developing a clinically-translatable subcutaneous adipose tissue regeneration strategy.

## Acknowledgements

Funding for this study was provided by the Canadian Institutes of Health Research (CIHR) (Operating Grant #119394), with infrastructure support from the Canadian Foundation of Innovation (CFI) and Ontario Research Fund-Research Infrastructure (ORF-RI) program. T.H. received scholarship support from the Ontario Graduate Scholarship (OGS) program and the Natural Sciences and Engineering Research Council of Canada (NSERC). The authors would like to acknowledge Dr. Aaron Grant, Dr. Robert Richards and Dr. Brian Evans for their clinical collaborations, as well as Dr. Laura Juignet, Dr. John Walker, Cody Brown, Anna Kornmuller and Kevin Robb for technical assistance.

---

**Supplementary Figure 1.**
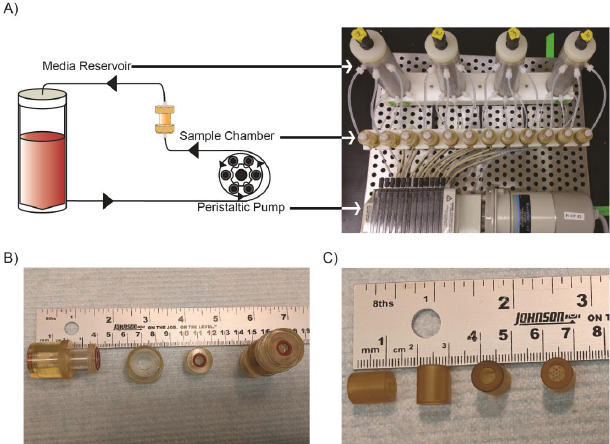
Schematic of the perfusion bioreactor system. A) Culture media is pumped unidirectionally via the peristaltic pump through the sample chamber before being recycled back into the media reservoir. B-C) The components of the sample chamber (B) and culture inserts (C).

**Supplementary Figure 2.**
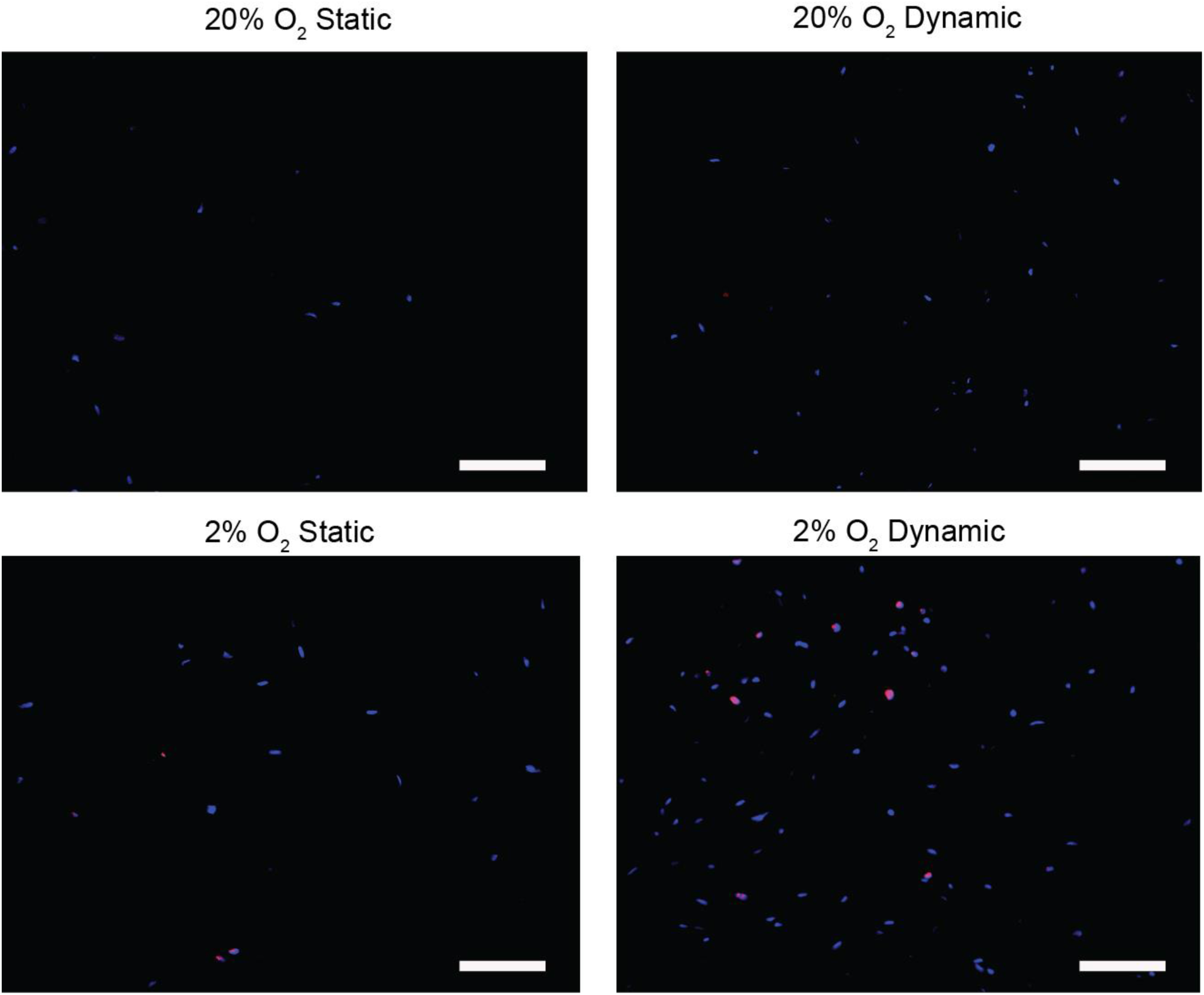
Representative Ki-67 staining in the central regions of the ASC-seeded DAT scaffolds cultured under static or dynamic conditions within the bioreactor for 14 days. Ki-67 expression is shown in red and DAPI counterstaining in blue. Scale bars represent 100 µm.

**Supplementary Figure 3.**
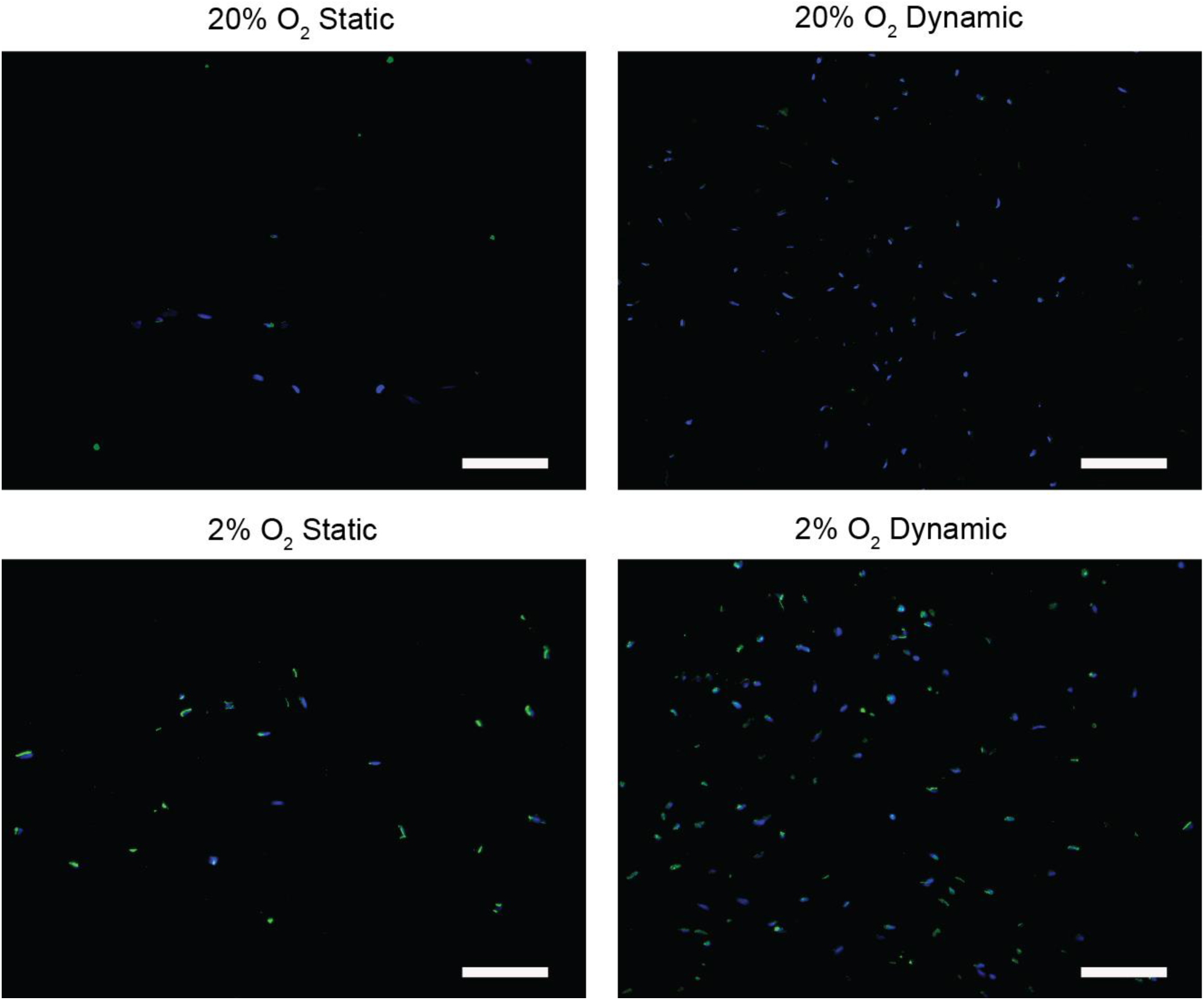
Representative HIF-1α staining in the central regions of the ASC-seeded DAT scaffolds cultured under static or dynamic conditions within the bioreactor for 14 days. HIF-1α^+^ cells are shown in green and DAPI counterstaining in blue. Scale bars represent 100 µm.

**Supplementary Figure 4.**
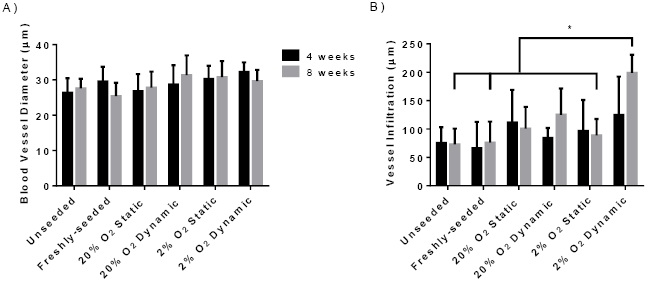
The average erythrocyte-containing blood vessel A) diameter and B) depth of infiltration in the DAT scaffolds following subcutaneous implantation in the *nu/nu* mouse model for 4 and 8 weeks. Error bars represent standard deviation (n=3 cross-sections/implant, N=6 implants per group). *=significant difference in the 2% O_2_ dynamic group as compared to the unseeded, freshly-seeded and 2% O_2_ static groups at 8 weeks (p<0.05).

**Supplementary Figure 5.**
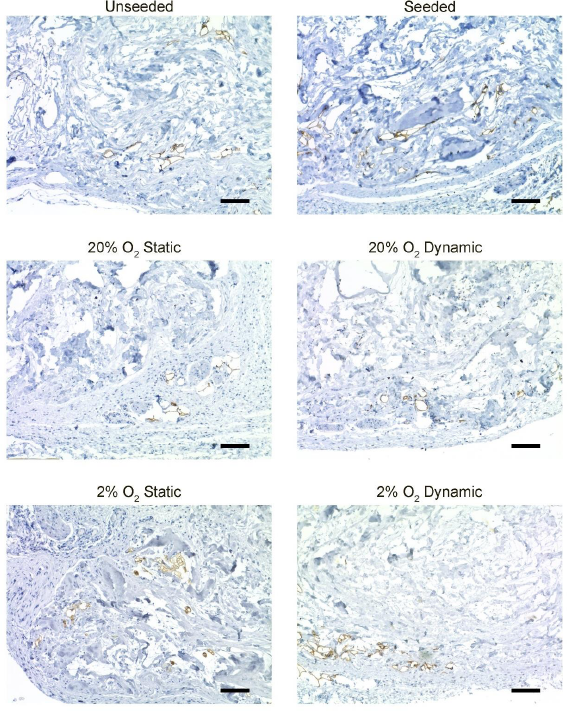
Representative images showing perilipin staining (brown) in the DAT scaffolds implanted subcutaneously in the *nu/nu* mouse model for 4 weeks. Scale bars represent 100 µm.

**Supplementary Figure 6.**
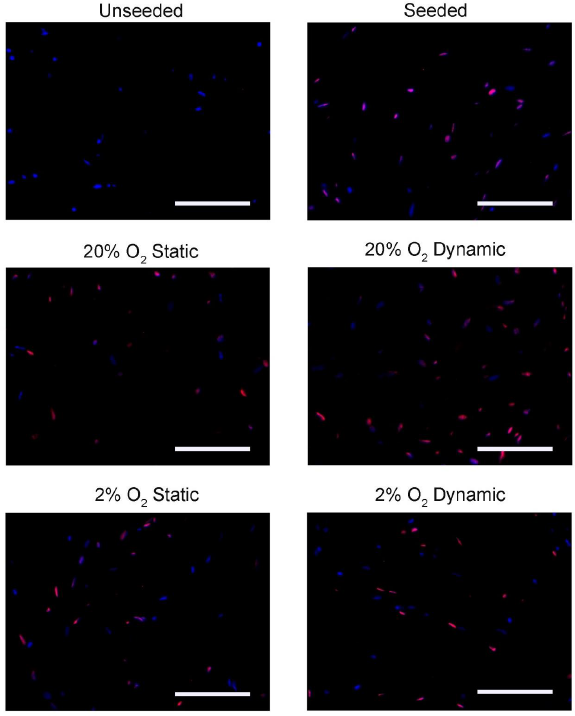
Representative immunostaining showing Ku80^+^ (red) human ASCs counterstained with DAPI (blue) in the DAT scaffolds that implanted subcutaneously in the *nu/nu* mouse model for 4 weeks. Scale bars represent 100 µm.

